# Impact of Comorbidities on SARS-CoV-2 Viral Entry-Related Genes

**DOI:** 10.1101/2020.05.26.117440

**Authors:** Joshua D. Breidenbach, Prabhatchandra Dube, Subhanwita Ghosh, Nikolai N. Modyanov, Deepak Malhotra, Lance D. Dworkin, Steven T. Haller, David J. Kennedy

## Abstract

Viral entry mechanisms for severe acute respiratory syndrome coronavirus 2 (SARS-CoV-2) are an important aspect of virulence. Proposed mechanisms involve host cell membrane-bound angiotensin-converting enzyme 2 (ACE2) and type II transmembrane serine proteases (TTSPs), such as transmembrane serine protease isoform 2 (TMPRSS2). The distribution of expression of these genes across cell types representing multiple organ systems in healthy individuals has been recently demonstrated. However, comorbidities such as diabetes and cardiovascular disease are highly prevalent in patients with Coronavirus Disease 2019 (COVID-19) and associated with worse outcomes. Whether these conditions contribute directly to SARS-CoV-2 virulence remain unclear. Here we show that the expression levels of *ACE2, TMPRSS2* and other viral entry-related genes are modulated in target organs of select disease states. In tissues such as heart, which normally express *ACE2* but minimal *TMPRSS2*, we found that *TMPRSS2* as well as other TTSPs are elevated in individuals with comorbidities vs healthy individuals. Additionally, we found increased expression of viral entry-related genes in the settings of hypertension, cancer or smoking across target organ systems. Our results demonstrate that common comorbidities may contribute directly to SARS-CoV-2 virulence and suggest new therapeutic targets to improve outcomes in vulnerable patient populations.

## Introduction

Comorbidities such as diabetes, chronic lung disease, and cardiovascular disease are highly prevalent in patients with COVID-19 and associated with worse outcomes (1, 2). However, whether these conditions contribute directly to SARS-CoV-2 virulence or simply worsen outcomes through independent mechanisms and reflect the general disease burden of the population remain unclear (2, 3). Furthermore, clinical and experimental evidence has demonstrated that in addition to the lungs, SARS-CoV-2 infection of other target organ systems such as heart, kidney, and blood may have important deleterious consequences which can potentially compromise organ function and compound disease burden in COVID-19 patients (4, 5).

The spike (S) protein of SARS-CoV and SARS-CoV-2 is a key facilitator for host cell entry through its binding to host cell membrane-bound ACE2 (6). Therefore, the impact of the modulation of *ACE2* and related renin-angiotensin-aldosterone system genes on COVID-19 has been an area of interest (3) and were included in the current study. Additionally, after binding, cleavage of the S protein is necessary for S protein-mediated membrane fusion which drives viral entry into host cells. This proteolytic activity may be cathepsin-L dependent and occur upon pH change in cellular endosomes, or may occur through the action of membrane bound serine proteases at the host cell membrane surface or within vesicles (7). Additionally, viral entry mechanisms have been proposed which involve cleavage of ACE2 itself by membrane bound serine proteases leading to increased viral entry (7). In fact, the importance of serine proteases in a viral entry mechanism may be emphasized by the success of serine protease inhibition in vitro (6). However, while this and current mechanistic studies have focused on the proteolytic activity of TMPRSS2 and human airway trypsin-like protease (HAT, also referred to as TMPRSS11D) additional TTSPs are hypothesized to have similar extracellular cleavage activity (8) and were included in the current study. Furthermore, the influence of comorbidities on these genes is unknown. Thus, in the current study we examined the influence of comorbidities on the expression of key renin-angiotensin-aldosterone system and protease genes which may prime the cell entry mechanisms for SARS-CoV-2 across various organ systems.

## Results and Discussion

In a recent thorough report, *ACE2* and *TMPRSS2* expression was mapped across various body sites in normal healthy tissue by single cell RNA sequencing (9). However, a more complete model of viral entry for SARS-CoV and SARS-CoV-2 describe a potential role for TMPRSS2, HAT, and potentially other TTSPs (7). Therefore, to confirm these recent findings and to better understand the expression patterns of other TTSPs in a healthy setting, expression data was analyzed for 27 different body sites from 3 healthy individuals (Figure 1). In healthy human tissues, there was a diverse transcription of renin-angiotensin-aldosterone system related genes (*ACE, ACE2*, and *AGTR1*) as well as proteases (*ADAM17, TMPRSS1*-*5, TMPRSSD, TMPRSSE*, and *TMPRSS15*) which may prime the cell entry mechanisms for SARS-CoV-2 across various organ systems. The results from our analysis are in agreement with previously reported expression patterns (9). Specifically, both surveys found high levels of *ACE2* and/or *TMPRSS2* expression in colon, heart, kidney, lung, and prostate. In addition, our analysis suggests salivary gland, small intestine, testis, thyroid, and trachea as sites of high expression.

**Figure 1.**
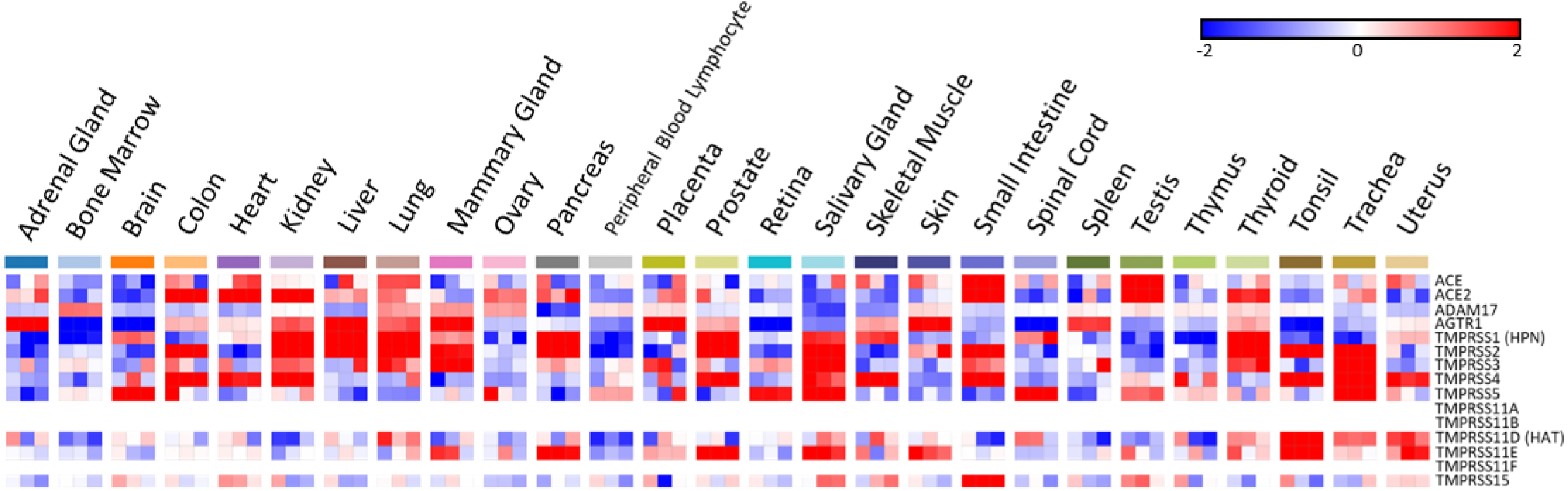
SARS-CoV-2 viral-entry gene expression across healthy human tissues. Expression of select genes related to SARS-CoV-2 viral-entry in healthy volunteers across 27 different body sites. Expression is displayed as logarithm to base 2 of the fold change (Log2FC), n=3. Heatmap is set to a scale of -2 (blue) to 2 (red). Blank cells indicate data missing from the DataSeries.

Additionally, we examined the impact of a variety of common comorbidities on the expression of these genes in select organ systems (pulmonary, renal, cardiac, and blood tissues) based on their relevance to infection and expression levels at baseline. Expression data displayed in each comorbidity heatmap is displayed from greatest (left) to least (right) expression of *ACE2*.

In pulmonary tissues (Figure 2A) we found the greatest increase in *ACE2* in cancer with substantial increases in nearly all of the TTSPs, which is consistent with their role in tumor cell proliferation, motility and invasion (8). Additionally, samples from patients with a history of smoking showed increases in *ACE2* in both small and large airways, consistent with recent findings (10). While expression levels in the context of pre-existing asthma appeared to be largely unaffected in pulmonary tissues on average, there were more pronounced increases in bronchial compared to nasal epithelium.

**Figure 2.**
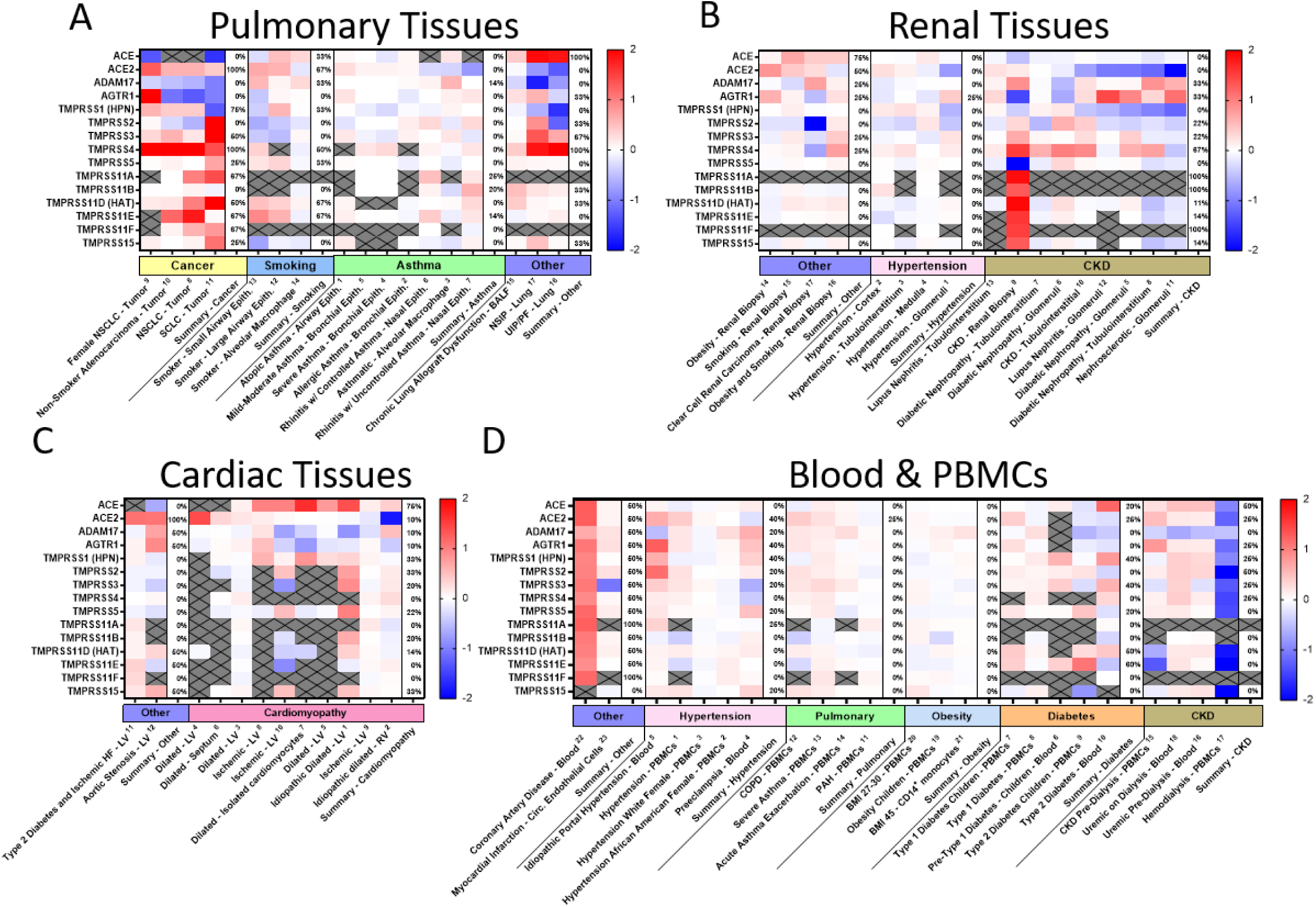
SARS-CoV-2 viral-entry gene expression across comorbidities. Expression of select SARS-CoV-2 viral-entry genes in pulmonary (A), renal (B), cardiac (C), and blood tissues (D) across 69 DataSets representing various common comorbidities. Values are displayed as logarithm to base 2 of the fold change (Log2FC) in comparison with respective unaffected control samples from each DataSet. Comorbidity groups and each DataSet within are sorted from greatest (left) to least (right) expression of *ACE2*. All heatmaps are set to a scale of -2 (blue) to 2 (red) and values beyond this range are shown as either 2 or -2. Grey crossed-out cells indicate data missing from the respective DataSet. Summary columns describe the percentage of the DataSets which demonstrated at least a 25% increase in expression of the respective gene within each comorbidity group shown. BALF = Bronchoalveolar lavage fluid; BMI = body mass index; CKD = chronic kidney disease; COPD = chronic obstructive pulmonary disease; HF = heart failure; LV = left ventricle; NSCLC = non-small cell lung carcinoma; NSIP = non-specific interstitial pneumonia; PBMCs = peripheral blood mononuclear cells; PAH = pulmonary arterial hypertension; RV = right ventricle; SCLC = small cell lung carcinoma; UIP/PF = usual interstitial pneumonia/idiopathic pulmonary fibrosis.

In renal tissues (Figure 2B), we found the greatest expression of *ACE2* in obesity. Similar to what has been seen in pulmonary tissues, a history of smoking or cancer associated with an increase in *ACE2* as well as slight increases in TTSPs in renal biopsy. Hypertensives had increases in *ACE2, TMPRSS1* and *TMPRSS4* in renal cortical and tubulointerstium, but not glomerular or medullar samples. Chronic kidney disease (CKD) resulted in the greatest diversity in modulation of these genes, however, with consistent increases in *TMRPSS4* in 67% of DataSets from both tubular and glomerular origin.

In cardiac tissues (Figure 2C), we found the greatest increases in *ACE2* in patients who had experienced heart failure with pre-existing diabetes or patients with aortic stenosis. While cardiomyopathies resulted in variable expression levels, increases in *ACE2* were found in left ventricle (LV) tissues while decreases were found in right ventricle (RV) tissues. On average, slight increases were found for many TTSPs with 33% of DataSets resulting in increases in *TMPRSS2*.

In blood (Figure 2D), remarkable increases in all selected genes were found in patients with coronary artery disease. Hypertension and chronic lung pathologies resulted in slight, but consistent increases in most of the selected genes including *ACE2* and many of the TTSPs. Specifically, in hypertension, increases were found in at least 20% of DataSets for *ACE2, ADAM17, AGTR1, TMPRSS1, TMPRSS2, TMPRSS3, TMPRSS5, TMPRSS11A*, and *TMPRSS15*. In contrast to the results seen in renal tissues, obesity did not appear to modulate the expression of any of these genes in circulating immune cells. Notably, increased expression levels were found in the context of type 1 diabetes, while decreases were apparent in type 2 diabetes in whole blood and peripheral blood mononuclear cells (PBMCs). Lastly, highly variable modulation was found in the context of CKD with or without hemodialysis with increases in *ACE, ACE2, AGTR1, TMPRSS1, TMPRSS2, TMPRSS3*, and *TMPRSS4* in at least 25% of DataSets.

Because interest in TTSPs seems to have blossomed only within the last decade, their appearances on micro array technologies and therefore their appearances in these data are limited. However, understanding the modulation of the expression of these genes across various organ systems and in the context of common comorbidities should provide us with a more complete understanding of the potential impact of these comorbidities on viral proliferation. Increases in the expression of these genes as suggested by this data in common comorbidities across tissues such as hypertension, cancer and a history of smoking may help to partially explain their association with higher morbidity and mortality in COVID-19. In other comorbidities, such as obesity, and diabetes or in those with tissue specificity such as in cardiomyopathies and chronic lung disease, the lack of consistent alteration across tissues may suggest a mechanism for tropism of the virus. To be sure, while SARS-CoV and SARS-CoV-2 virulence is driven by factors outside of viral entry, it may be important to understand the influence of the varied expression of these genes among target organ systems and across comorbidities as demonstrated.

In conclusion, expression levels of SARS-CoV-2 viral-entry related genes in patients suffering from common comorbidities such as hypertension, cancer, a history of smoking, obesity, diabetes, cardiomyopathies, or chronic lung or kidney disease may be increased in target organ systems and be capable of directly contributing to infection. This represents an important step in designing effective therapeutic and preventative strategies to improve outcomes in vulnerable populations.

## Methods

In the current study, we examined the expression levels of SARS-CoV-2 entry-related genes in target organ systems including pulmonary, renal, cardiac, and PBMCs from 2032 patients across a variety of common comorbidities including hypertension (n=94 hypertensives vs. n=61 normotensives), diabetes (n=131 diabetic vs. n=101 normoglycemic), obesity (n=56 obese vs. n=58 healthy weight), chronic lung disease (n=200 asthmatic vs. n=106 non-asthmatic and Chronic Obstructive Pulmonary Disease (COPD) n=94 vs. healthy tissue n=42) and cardiovascular disease (ischemic n=36 and dilated n=91 cardiomyopathy vs. healthy tissue n=69), as well as other common pathologies such as CKD and cancer. Differential gene expression was curated from genetic data deposited in the National Center for Biotechnology Information (NCBI), U.S. National Library of Medicine, Gene Expression Omnibus (GEO) DataSets and the European Molecular Biology Laboratory (EMBL), European Bioinformatics Institute (EBI). Exhaustive queries in these databases in the form of “[tissue] and [hypertension, diabetes, obesity, smoking, asthma]” were performed using the NCBI and iLINCS website (ilincs.org) and all hits were considered. DataSeries not containing disease state vs. healthy controls for the tissues concerned were excluded and common off-target hits such as cancer were included. Differential Expression analysis was performed using the GEO2R (NCBI) interactive web tool, and the iLINCS integrative web platform for the analysis of the Library of Integrated Network-Based Cellular Signatures (LINCS). The expression values in healthy tissues are reported as logarithm to base 2 of the fold change (Log2FC) as described in the GEO DataSet (GDS3113) and visualized by Morpheus (Broad Institute). For comorbidity disease condition data, expression levels are described in log2FC in relation to each unaffected control group specific to each DataSeries and heatmaps were generated in GraphPad Prism (GraphPad Software Inc.). All log2FC values outside of the -2 to 2 range are shown as either -2 or 2. DataSeries in this analysis were as follows with GSEXXXX referring to NCBI-GEO and E-XXXX-XXXX referring to EMBL-EBI:

### Pulmonary Tissues

(1, GSE18965; 2, GSE41649; 3, GSE2125; 4, E-GEOD-63142; 5, E-GEOD-63142; 6, E-GEOD-19187; 7, E-GEOD-19187; 8, E-GEOD-44077; 9, GSE19804; 10, E-GEOD-43458; 11, E-GEOD-60052; 12, GSE5057; 13, GSE3320; 14, GSE2125; 15, E-MTAB-6040; 16, GSE21411; 17, GSE21411);

### Renal Tissues

(1, GSE104948; 2, GSE28345; 3, GSE104954; 4, GSE28360; 5, GSE104948; 6, GSE30528; 7, GSE104954; 8, GSE30529; 9, GSE66494; 10, GSE12682; 11, GSE20602; 12, GSE32591; 13, GSE32591; 14, GSE46699; 15, GSE46699; 16, GSE46699; 17, GSE46699);

### Cardiac Tissues

(1, GSE1145; 2, GSE67492; 3, GSE42955; 4, GSE3585; 5, GSE116250; 6, GSE3586; 7, GSE120836; 8, GSE116250; 9, GSE42955; 10, GSE1145; 11, GSE26887; 12, GSE10161;

### Blood and PBMCs

(1, GSE24752; 2, GSE75360; 3, GSE75360; 4, GSE48424; 5, GSE69601; 6, E-TABM-666; 7, GSE9006; 8, GSE55098; 9, GSE9006; 10, GSE23561; 11, GSE131793; 12, GSE42057; 13, GSE59019; 14, GSE16032; 15, GSE15072; 16, GSE37171; 17, GSE15072; 18, GSE37171; 19, GSE87493; 20, GSE69039; 21, GSE32575; 22, GSE23561; 23, GSE66360).

## Author Contributions

JDB, PD, NNM, LDD, DM, STH, and DJK conceptualized the study. JDB, PD, and SG acquired the data. JDB, PD, NNM, STH, and DJK analyzed the data. JDB wrote the original draft of the manuscript. All authors reviewed and edited the manuscript. JDB, PD, STH, and DJK were responsible for visualization. DM, STH, and DJK acquired funding.

## Acknowledgments

This work was supported by the National Institutes of Health (HL-137004) the David and Helen Boone Foundation Research Fund, the University of Toledo Women and Philanthropy Genetic Analysis Instrumentation Center, and the University of Toledo Medical Research Society. The authors would like to thank Rick Gerasimiak, Steve Acton, and the University of Toledo Department of Neurosciences for their guidance.

